# RNA-TorsionBERT: leveraging language models for RNA 3D torsion angles prediction

**DOI:** 10.1101/2024.06.06.597803

**Authors:** Clément Bernard, Guillaume Postic, Sahar Ghannay, Fariza Tahi

**Affiliations:** Université Paris Saclay, Univ Evry, IBISC, 91020, Evry-Courcouronnes, France; LISN - CNRS/Université Paris-Saclay, France, 91400, Orsay, France

**Keywords:** RNA 3D structure, Torsional angles, Language models, Scoring function

## Abstract

Predicting the 3D structure of RNA is an ongoing challenge that has yet to be completely addressed despite continuous advancements. RNA 3D structures rely on distances between residues and base interactions but also backbone torsional angles. Knowing the torsional angles for each residue could help reconstruct its global folding, which is what we tackle in this work. This paper presents a novel approach for directly predicting RNA torsional angles from raw sequence data. Our method draws inspiration from the successful application of language models in various domains and adapts them to RNA. We have developed a language-based model, RNA-TorsionBERT, incorporating better sequential interactions for predicting RNA torsional and pseudo-torsional angles from the sequence only. Through extensive benchmarking, we demonstrate that our method improves the prediction of torsional angles compared to state-of-the-art methods. In addition, by using our predictive model, we have inferred a torsion angle-dependent scoring function, called TB-MCQ, that replaces the true reference angles by our model prediction. We show that it accurately evaluates the quality of near-native predicted structures, in terms of RNA backbone torsion angle values. Our work demonstrates promising results, suggesting the potential utility of language models in advancing RNA 3D structure prediction. Source code is freely available on the EvryRNA platform: https://evryrna.ibisc.univevry.fr/evryrna/RNA-TorsionBERT.

## Introduction

RNA is a macromolecule that plays various biological functions in organisms. Similarly to proteins, the biological function of an RNA may be directly linked to its 3D structure. Experimental methods such as NMR, X-ray crystallography, or cryo-EM can determine the 3D structure of RNAs, but they remain tedious in cost and time. Computational methods have been developed for predicting the 3D structure from the sequence, with three different approaches: *ab initio*, template-based and deep learning-based (1). Currently, no existing method matches the performance of AlphaFold 2 for proteins (2), as shown with the last results on the CASP-RNA challenge (3). Reaching AlphaFold’s (2) level of accuracy is a long shot, notably due to the lack of data (4). Very recently, the release of AlphaFold 3 (5) has extended its predictions to a wide range of molecules like DNA, ligand, ion and RNA, but the results remain limited for RNA (5, 6).

RNA can adopt various secondary motifs, along with a wide range of complex interactions that contribute to its 3D structure. Research efforts have focused on classifying both the canonical and non-canonical pairs, further supported by the description of the backbone conformation (7). Unlike proteins, RNA backbone structures are defined by eight torsional angles, the natural manifold of all these dihedral angles combined being an 8-dimensional hypertorus, which presents a significant challenge both statistically and computationally. Pyle and colleagues have shown that they can be approximated by two pseudo-torsional angles (see Figure 1) (8). Understanding these torsional angles is crucial for comprehending the 3D structures of RNA, which in turn could aid in predicting their folding.

**Fig. 1.**
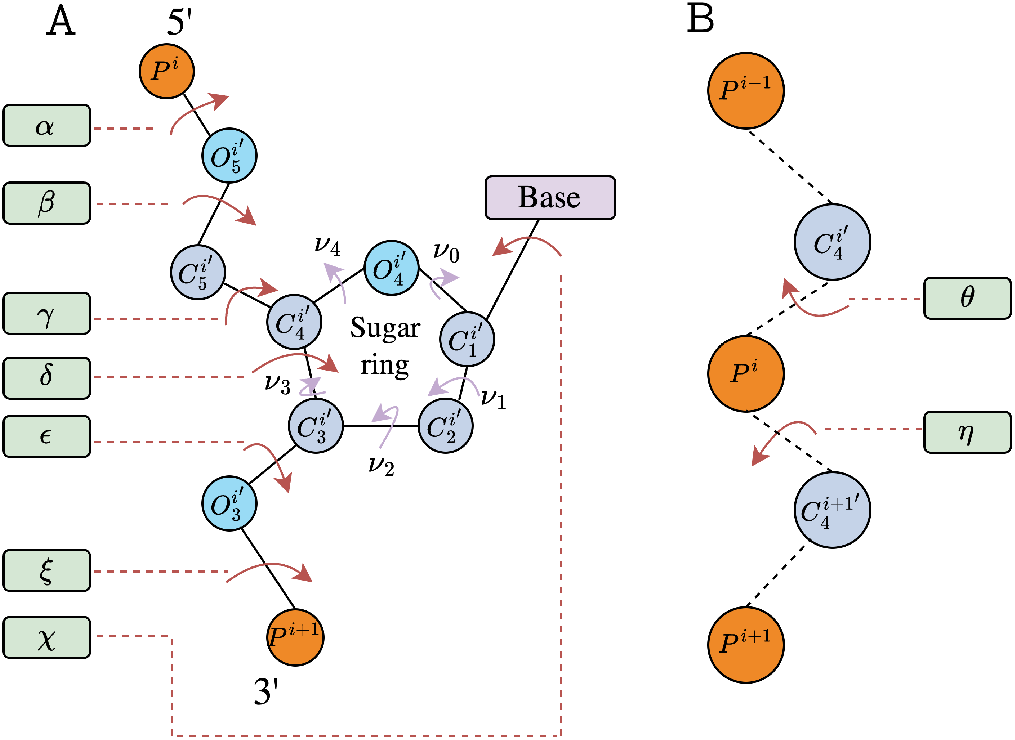
The eight RNA torsional angles and the two pseudo-torsional angles. (A) RNA backbone torsional angles. The angles are defined around 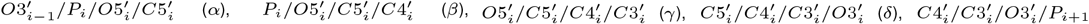 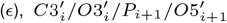 (*ζ*) and the rotation of the base relative to the sugar 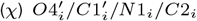 for pyrimidines and 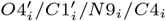 for purines. The ribose ring angles are defined as 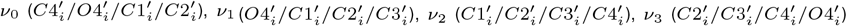 and 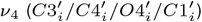 The ribose ring is usually described by a single sugar pucker pseudorotation phase 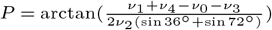 (B) RNA pseudo-torsional angles. *η* is defined around 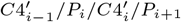 and *θ* around 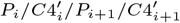

Current predictive methods for RNA 3D structure prediction do not always integrate torsional angles, missing important features to comprehend its folding. One work (9) has focused on constructing libraries of RNA conformers with torsional angles. It has been used for RNAfitme (10), which allows editing and refining predicted RNA 3D structures. Another work has been done to predict exclusively torsional angles from RNA sequence, SPOT-RNA-1D (11), using a residual convolutional neural network.

In this work, we aim to leverage language models to better apprehend the prediction of RNA torsional angles from its sequence. Indeed, works have been proposed through the years to work on biological sequences, inspired by the success of language models like BERT (12). Its adaptation for RNA (13) or DNA (14) shows promising results which could be leveraged for other RNA structural features prediction.

Another interest in torsional angles for RNA 3D structures is for quality assessment. Without the help of reference structures, scoring functions have been developed to assess structural quality. These methods can be knowledge-based (15– 17) using statistical potentials or deep learning (18). The knowledge-based scoring functions consider RNA structural features as inputs like pairwise distances (15, 16, 19) or with the help of torsional angles (17). We propose here a new scoring function based on the extension of our model to predict RNA torsional angles. This scoring function allows us to assess structural quality in torsional angle space.

This paper is organized as follows: we describe our contributions in two separate points. Each section is divided into two parts: one for the work on torsional angle prediction and the other for our proposed scoring function. We detail our experiments for the torsional angles prediction and the structural quality assessment tasks before discussing our approaches’ results and limitations. We then conclude by discussing the scope of our contributions. The results and the code of our RNA-TorsionBERT and TB-MCQ are easily reproducible and freely available on the EvryRNA platform: https://evryrna.ibisc.univevry.fr/evryrna/RNA-TorsionBERT.

## Methods

This section presents our model for predicting RNA torsional angles and then the scoring function derived from our model.

### Torsional angles prediction

#### RNA-TorsionBERT approach

Current methods that use sequence as inputs for RNA-related approaches only represent sequences as one-hot-encoding vectors. This representation may be too sparse to consider sequential interactions well. This encoding is usually associated with a convolutional neural network, which is commonly limited by longrange interactions. A solution could be using attention mechanisms. Attention-based architecture nonetheless requires a huge amount of data to train well, which is not the case for RNA 3D structure data. To counter this problem, we can use models pre-trained on a large amount of unlabeled data. This could bring a better input representation of the raw sequence, which could then be fine-tuned to specific tasks. These pretrained models could input either RNA or DNA sequences. Recent advances in language models started with BERT (12), where the model was pre-trained on masking and next-sentence prediction tasks before being fine-tuned on diverse specific language tasks. DNA or RNA can be seen as a sequence of nucleotides, where their interactions have a biological meaning. Therefore, methods have been adapted from BERT to develop a language-based architecture for either RNA or DNA. The aim is to reproduce the success of language comprehension for another language. As the size of the vocabulary is different, modifications should be made to fit the current language. An example for DNA is DNABERT (14), where the training process was updated compared to the original BERT by removing the next sentence prediction and taking *K*-mers as inputs (contiguous sequence of k nucleotides). It was trained on human-genome data. An example of adaptation of BERT for RNA is called RNABERT (13). It is a six-transformer layer pre-trained on two tasks: masking and structural alignment learning (SAL). RNABERT was trained on 76,237 human-derived small ncRNAs. Other methods have been adapted to RNA language but uses MSA as inputs like RNA-MSM (20). Nonetheless, they require multiple sequence alignment (MSA) as inputs, which restricts the use for RNAs. Indeed, there are a numerous amount of unknown structures (21), and MSA will restrict the adaptation to future unseen families. In this article, we decided to only consider sequences as inputs, and so for the language models.

The aim of our method is, given a pre-trained language model (DNABERT or RNABERT), to adapt its neuronal weights to predict RNA torsional and pseudo-torsional angles from the sequence. We have added layers to adapt the methods to our multi-token label regression task. Each token in the input would have 28 labels: two values (sine and cosine) for each of the eight torsional angles (the phase *P* being represented by its five ribose ring angles) and two pseudo-torsional angles. The use of pre-trained embedding would help the model not to start from scratch and update the learned attention layers for RNA structural features.

#### RNA-TorsionBERT architecture

The architecture of our method, when based on DNABERT, is described in Figure 2 (illustrated with 3-mers). An input sequence of size L is tokenized and then fed to the network with token and positional embeddings. The tokenization process usually adds specific tokens (like the *CLS* and *PAD* tokens). As DNABERT could input a maximum of 512 tokens, we set the maximum sequence length to 512 nucleotides. The last hidden state is set to be 768 by the original DNABERT architecture. We then apply extra layers to map the hidden state outputs to the desired final output dimension (Lx28). These extra layers comprise layer normalisation, a linear layer (from 768 to 1024), a GELU activation, another linear layer (1024 to 28), and a Tanh final activation. The final output layer is of size 28 because it outputs a sine and a cosine for the eight torsional (*α, β, γ, δ, E, ζ, χ* and the phase *P* being predicted through the five ribose ring angles *ν*_0_, *ν*_1_, *ν*_2_, *ν*_3_ and *ν*_4_) and two pseudo-torsional angles (*η* and *θ*). It allows the relief of the periodicity of the different angles. The Tanh activation maps the outputs to the cosine and sine range (between −1 and +1), which is then converted into angle predictions using the formula 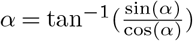 (adaptable for the other angles). Details on the training process are in the Supplementary file.

**Fig. 2.**
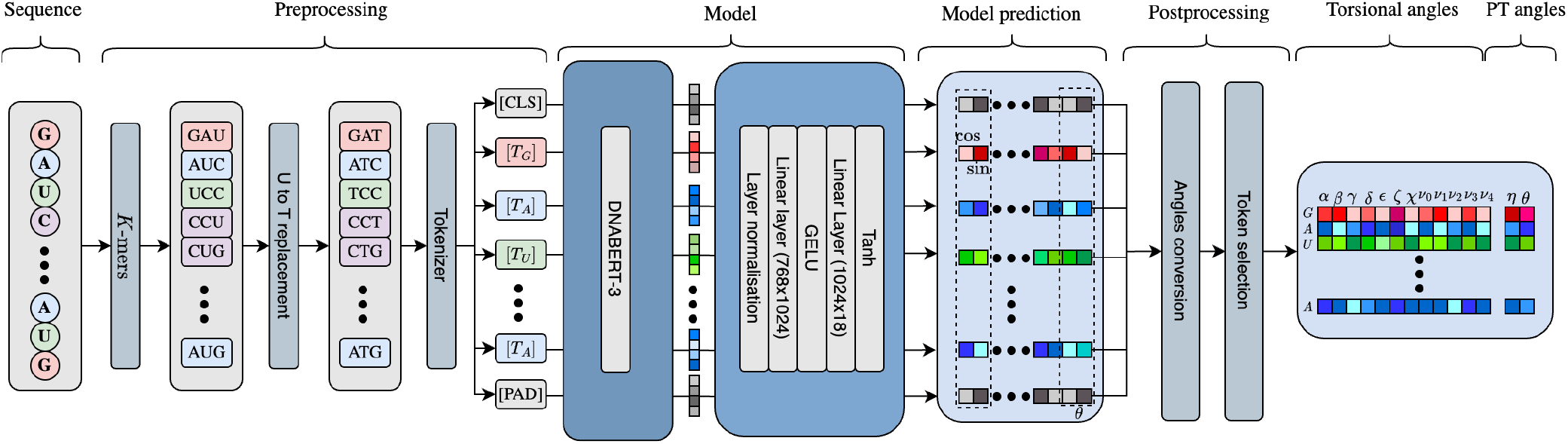
Schema of the proposed language model architecture for torsional and pseudo-torsional angles prediction. Given an RNA sequence, mapping is applied to each sequence’s nucleotide into a token with embeddings from the language model. *CLS* and *PAD* tokens are added to the sequence tokens. We convert the Uracil (U) with its equivalent in DNA: Thymine (T) for DNABERT model. Then, the language model will output hidden states with a representation of each token. This is fed into extra layers before entering a last Tanh activation to have the cosine and sine per angle. A postprocessing is required to convert back to angles (and pseudo-torsional (PT) angles) from the sine and cosine.

### Model quality assessment based on torsional angles

#### Torsional-based quality assessment metrics

Existing metrics have been developed to assess the quality of predicted RNA 3D structures with access to a reference. The most famous one is the RMSD (root-mean-square deviation), which assesses the general folding of structures. Other metrics have been developed and adapted from proteins (22, 23). Some specific metrics have also been designed to consider RNA specificities (24). Only two metrics are torsional-angles-based: the MCQ (25) (mean of circular quantities), and the Longest Continuous Segment in Torsion Angle space (LCS-TA) (26). The MCQ computes the deviation in angle space without any superposition of structures and complements other existing metrics. LCS-TA computes the longest number of continuous residues with an MCQ below a threshold (usually 10°, 15°, 20° and 25°). It is also a superposition-independent metric.

In SPOT-RNA-1D (11), the authors introduced the mean-average error (MAE) metric to assess the performance of their method SPOT-RNA-1D in the prediction of torsional and pseudo-torsional angles. Nonetheless, the MAE is an arithmetic mean and is not designed for angles.

To compute deviation for circular quantities, we use the mean of circular quantities (MCQ) (25). We don’t consider the LCS-TA as it is more expensive to compute, and the MCQ is more widely used in RNA-Puzzles (27–30). We define the set of angles for the torsional angles as *T* = {*α, β, γ, δ, E, ζ, P, χ*} and for pseudo-torsional angles *PT* = {*η, θ*}. Following the notation in (25), for a given structure *S* of *L* residues, let’s note *t*_*i*,*j*_ the torsional angle of type *j* of the residue at position *i*. We denote the difference between two structures *S* and *S*^*l*^ as the MCQ(S,S’), defined by:

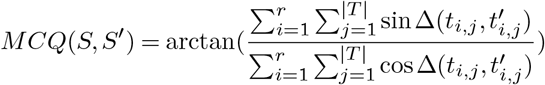

where *r* is the number of residues in *S* ∩ *S*^*l*^ and with:

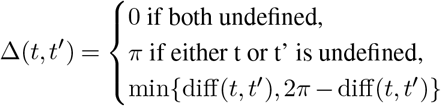

and:

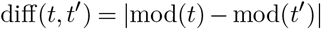

The difference aims to consider periodicity of 2*π* with

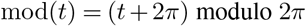

To have more details on the performances for a specific angle, we define the MCQ for a specific type of angle *j*:

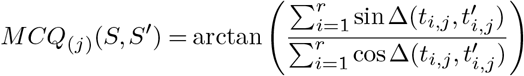

We extend the formulation for pseudo-torsion angles by just changing the set of angles used, and we name it *MCQ*_*PT*_.

#### Quality assessment scoring functions

Quality assessment of RNA 3D structures requires two structures, with one being the experimentally solved structure. Having this reference is a strong asset that is hardly possible in practice. To rank near-native structures without a reference, scoring functions have been developed, adapting free-energy (15–17). Other methods employ deep learning approaches like ARES (18).

To discriminate near-native structures in the torsional space, we have derived a scoring function from our RNA-TorsionBERT model. First, we have replicated a quality assessment metric that uses torsional angles features: the mean of circular quantities (MCQ) (25). Then, we replaced the true torsional angles with the predicted angles from our model to compute the MCQ over the near-native structure. Therefore, the MCQ computation compares the prediction of our model angles with the angles from the predicted non-native structures. This MCQ now becomes a scoring function, as it only takes as input a structure without any known native structure. We named this scoring function TB-MCQ for TorsionBERT-MCQ. Figure 3 shows the architecture of TB-MCQ. Given a structure, we extract the torsional angles and the sequence. The sequence is then pre-processed by RNA-TorsionBERT, and the inference gives predicted angles. Then, we compute the MCQ to finally output a structural quality measure for an RNA 3D structure.

**Fig. 3.**
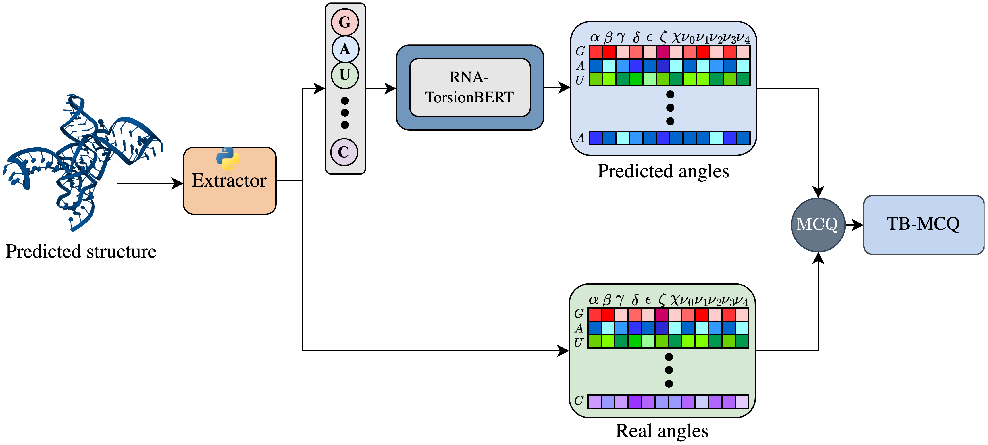
TB-MCQ scoring function computation. Given a predicted RNA 3D structure, we extract the sequence and calculates the different torsional angles. The sequence is fed to our RNA-TorsionBERT model to predict torsional angles. The scoring function is computed by taking the MCQ between the predicted angles and the real angles.

## Results and Discussion

### Results on Torsional Angles prediction

We present here the different experiments for the torsional angles prediction task. We used the MCQ presented above as a criterion to assess our model performances. We mainly focus on the results for torsional angles, while the results of *MCQ*_*PT*_ for pseudo-torsional angles are available in the Supplementary file.

#### Datasets

To validate the performances of torsional angle prediction models, we used different datasets of native structures:

### Training

we downloaded each available PDB structure and removed the structures from the non-redundant Validation and Test sets presented below. We also ensure the structures from this dataset have a sequence similarity below 80% compared to the other used datasets. We considered only the structures of a maximum sequence length of 512 (DNABERT can only input 512 tokens). The final set is composed of 4,267 structures with sequences from 11 to 508 nucleotides.

### Validation

we used the validation structures from SPOT-RNA-1D (11). It contains 29 structures with sequences between 33 and 288 nucleotides.

### Test

we combined two well-known test sets: RNA-Puzzles (31) and CASP-RNA (3). We combined both of these datasets as a whole Test set to assess the robustness of our model. It leads to a Test set of 34 structures (22 from single-stranded RNA of RNA-Puzzles and 12 from CASP-RNA), with sequences from 27 nucleotides to 512 (we cropped the RNA of PDB ID R1138 (720nt) to 512 nucleotides).

The distribution of the eight torsional angles and the two pseudo-torsional angles is given in Figure 4. As the pseudorotation phase *P* is defined with the five ribose ring angles, their distributions are shown in Figure S1 of the Supplementary file. These distributions are similar for the three datasets, meaning the learned distribution from the training set could allow good generalisation for the model.

**Fig. 4.**
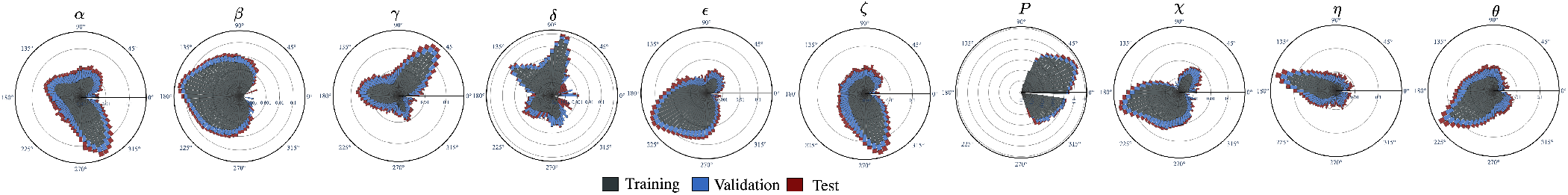
Polar distribution of the eight torsional angles (*α, β, γ, δ, ζ, P*, and *χ*) and the two pseudo-torsional angles (*η* and *θ*) for the Training, Validation and Test datasets. For each angle, the logarithm of the normalized count is depicted.

#### Language model selection

Each language model has a different format of inputs (*K*-mers for DNABERT and single nucleotides for RNABERT), we had to select the best tokenization of our RNA sequences. We also had to decide which pre-trained model was the best for our task. Therefore, we trained the same DNABERT model with the different values of K (3, 4, 5 or 6) and RNABERT on the Training set and observed the performances on the Validation set.

The results are shown in Table 1 on the Validation set. In terms of *K*-mers, DNABERT trained on 3-mers has better results (MCQ of 19.0) than the other *K*-mers and RNABERT, even if it does not outperform them for each torsional angle. The results for the pseudorotation phase *P* show that the model does not change the prediction for this angle. RNABERT only outperforms the other methods for the *P* angle, which does not lead to any significant conclusion for the selection of this model. We observe that for some angles (*β, P* and *χ*), the choice of models does not have an impact on the performances. DNABERT with 3-mers outperforms RNABERT, which does not input *K*-mers. This result remains surprising as we could have thought that RNABERT, as pre-trained specifically on RNA data, could have done better than the DNABERT model. This difference might be explained by the *K*-mers representation that is used by DNABERT compared to RNABERT, where the size of the vocabulary is extended, and thus a finer representation of the inputs is embedded. This could help the model learn a higher number of interactions and be more adaptable for other tasks. What could also explain the difference in performances is the size of the model: DNABERT has a size of around 328MB, whereas RNABERT has around 2MB.

**Table 1.** MCQ for each torsional angle and using all the torsional angles for the Validation set for DNABERT (3,4, 5 or 6-mers) and RNABERT.

|  | RNABERT |  | DNABERT |  |  |  |
| --- | --- | --- | --- | --- | --- | --- |
|  | - | 3-mer | 4-mer | 5-mer | 6-mer |  |
| $MCQ(\alpha)$ | 32.3 | **31.0** | 31.8 | 33.9 | 36.3 | |
| $MCQ(\beta)$ | **17.6** | **17.6** | **17.6** | 17.8 | 17.8 | |
| $MCQ(\gamma)$ | 26.3 | **22.8** | 26.5 | 26.7 | 28.1 | |
| $MCQ(\delta)$ | 14.4 | **12.1** | 15.6 | 16.7 | 17.3 | |
| $MCQ(\epsilon)$ | 16.1 | **15.8** | 16.0 | 16.0 | 16.0 | |
| $MCQ(\zeta)$ | 22.5 | 21.7 | **21.6** | 21.8 | 22.2 | |
| $MCQ(P)$ | **8.6** | 8.7 | 8.8 | 8.7 | 8.7 | |
| $MCQ(\chi)$ | **18.2** | **18.2** | 18.3 | 18.5 | 18.7 | |
| $MCQ$ | 20.2 | **19.0** | 20.2 | 20.7 | 21.4 | |

From now on, we name RNA-TorsionBERT (for RNA torsional BERT) the DNABERT with 3-mers.

#### Performances

We present here the prediction results obtained by our method RNA-TorsionBERT on the Test set (presented above) compared to the state-of-the-art approach SPOT-RNA-1D (11), which is the only method that predicts RNA torsional angles from the sequence only. We reproduced the architecture of SPOT-RNA-1D (because we only had the code to do inference) and trained it with the exact same data as RNA-TorsionBERT. We also included the results for the inferred angles from methods benchmarked in State-of-the-RNArt (1). The methods benchmarked included either *ab initio* with IsRNA (32) and RNAJP (35), or template-based with RNAComposer (34), Vfold-Pipeline (33), MC-Sym (37) and 3dRNA (36). We also include three deep learning methods: trRosettaRNA (38) and RhoFold (39) and the newly AlphaFold 3 (5). We report the MCQ per angle on the Test Set in Table 2. *MCQ*_*PT*_ (pseudo-torsional) results are available in Table S1 of the Supplementary file. Our RNA-TorsionBERT model has better performances than SPOT-RNA-1D for every angle. It has an average MCQ of 17.4 compared to 19.4 for SPOT-RNA-1D. The *MCQ* improvement over SPOT-RNA-1D ranges between 0.2° (for *E*) and 4.3° (for *δ*). It also outperforms the angles inferred from state-of-the-art methods for RNA 3D structure prediction, including the last published method, AlphaFold 3 (5). Nonetheless, the performances compared to AlphaFold 3 remains close. RNA-TorsionBERT does not outperform it for every angle. trRosettaRNA and RhoFold, two deep learning methods, have the worst MCQ compared to *ab initio* and template-based approaches. It can be explained by the use of physics in *ab initio* and template-based methods that are inferred in the torsional angles. The use of deep learning approaches might have the counterpart to not include physics enough, except for AlphaFold 3. Deep learning methods, while having the best overall results, as shown in the benchmark done in State-of-the-RNArt (1), remain limited in torsional angle predictions.

**Table 2.** MCQ per torsional angle and for all torsional angles over the Test set for RNA-TorsionBERT compared to SPOT-RNA-1D. We also include inferred torsional angles from state-of-the-art methods that predict RNA 3D structures from State-of-the-RNArt (1). Methods are either deep learning (DL), ab initio (AB) or template-based (TP).

| | Type | $MCQ_{(\alpha)}$ | $MCQ_{(\beta)}$ | $MCQ_{(\gamma)}$ | $MCQ_{(\delta)}$ | $MCQ_{(\epsilon)}$ | $MCQ_{(\zeta)}$ | $MCQ_{(P)}$ | $MCQ_{(X)}$ | MCQ |
| --- | --- | --- | --- | --- | --- | --- | --- | --- | --- | --- |
| RNA-TorsionBERT | DL | 29.9 | **19.0** | **23.7** | 9.5 | 15.3 | 19.1 | **8.4** | **12.2** | **17.4** |
| SPOT-RNA-1D (11) | DL | 32.5 | 19.6 | 26.6 | 13.7 | 15.5 | 20.2 | 9.8 | 13.2 | 19.4 |
| AlphaFold3 (5) | DL | **29.8** | 19.9 | 23.9 | **8.8** | **15.2** | **18.8** | 8.8 | 14.1 | 17.8 |
| IsRNA1 (32) | AB | 41.9 | 23.8 | 33.5 | 12.5 | 22.8 | 31.3 | 18.9 | 17.5 | 24.9 |
| Vfold-Pipeline (33) | TP | 41.3 | 24.1 | 32.3 | 14.7 | 23.4 | 29.3 | 17.3 | 20.6 | 25.3 |
| RNAComposer (34) | TP | 43.6 | 27.5 | 38.8 | 13.3 | 21.4 | 27.4 | 16.4 | 20.6 | 25.9 |
| RNAJP (35) | AB | 41.3 | 28.0 | 33.3 | 14.0 | 24.4 | 32.1 | 11.2 | 20.7 | 26.6 |
| 3dRNA (36) | TP | 50.4 | 31.9 | 42.3 | 21.4 | 31.0 | 36.1 | 24.2 | 23.2 | 32.5 |
| MC-Sym (37) | TP | 66.5 | 26.0 | 57.9 | 27.8 | 22.1 | 39.6 | 17.5 | 23.4 | 36.0 |
| trRosettaRNA (38) | DL | 59.1 | 33.8 | 60.2 | 21.9 | 27.9 | 41.1 | 28.3 | 55.4 | 40.4 |
| RhoFold (39) | DL | 91.4 | 61.3 | 67.4 | 48.1 | 45.0 | 53.6 | 46.7 | 32.3 | 54.8 |

#### Results according to sequence length

To study more in details the performances based on the RNA length, we report in Figure 5 the MCQ obtained by our method, SPOT-RNA-1D and AlphaFold 3 depending on the sequence length for the Test set. We can see our method outperforms SPOT-RNA-1D for each of the sequence slot. AlphaFold 3 has lower MCQ for structures with sequences between 75 and 175nt. For sequences higher than 200 nucleotides, our method demonstrates superior performances compared to both SPOT-RNA-1D and AlphaFold 3, showing the interest for long range sequences. Results for the *MCQ*_*PT*_ are shown in Figure S2 of the Supplementary file.

**Fig. 5.**
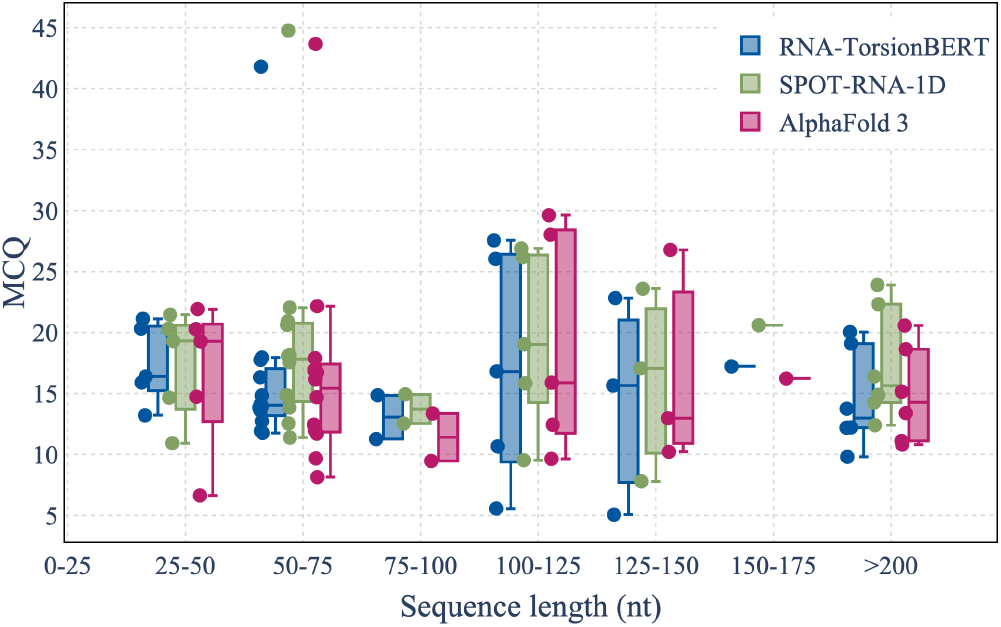
MCQ depending on sequence length (with a window of 25nt from 25nt to 200nt) for RNA-TorsionBERT, SPOT-RNA-1D and AlphaFold 3 for the Test set.

#### Results according to secondary structure motifs

We report the results of MCQ for three types of secondary structure motifs (single-stranded, loop and stem) averaged over the Test set for RNA-TorsionBERT, AlphaFold 3 and SPOT-RNA-1D in Table 3. We observe that our method delivers improved performances for each secondary structure motif (extracted from RNApdbee (40)). It has an overall MCQ higher for single-stranded than stem motifs. This behaviour is also similar to SPOT-RNA-1D and AlphaFold 3, which could be explained by the fact that stem and loop motifs are easier to predict than single-stranded motifs (and so are the base pairings). Details on the results for pseudo-torsional angles are available in Table S2 of the Supplementary file.

**Table 3.**
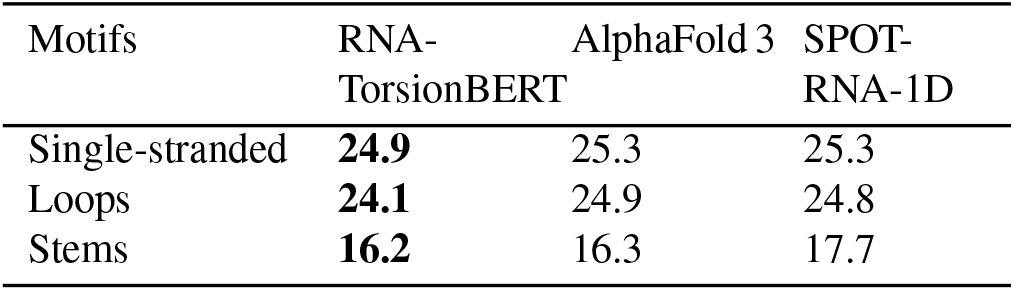
MCQ for torsional angles averaged over the Test set for RNA-TorsionBERT, AlphaFold 3 and SPOT-RNA-1D for three secondary structure motifs: singlestranded, loops and stems. Motifs are extracted using RNApdbee (40).

| Motifs | RNA-TorsionBERT | AlphaFold 3 | SPOT-RNA-1D |
| --- | --- | --- | --- |
| Single-stranded | **24.9** | 25.3 | 25.3 |
| Loops | **24.1** | 24.9 | 24.8 |
| Stems | **16.2** | 16.3 | 17.7 |

#### Results according to RNA types

In CASP-RNA, structures can be described as either natural with or without homologs, or synthetic RNAs. To further study the different cases where our approach is better than existing tools, we report the results for the natural (with or without homologs) and synthetic RNAs in Table 4. Our method outperforms AlphaFold 3 and SPOT-RNA-1D for natural RNAs, with the largest gap for RNA without homologs. This could be explained by the reliability of AlphaFold 3 on multiple sequence alignment, and, thus, on the availability and quality of homologs for the prediction. AlphaFold 3 has better performances for synthetic RNAs. More details on the results for different RNA families on RNA-Puzzles are available in Table S3 of the Supplementary file. Examples of structures are provided in Figure S3 of the Supplementary file.

**Table 4.** MCQ per RNA type for the CASP-RNA dataset for RNA-TorsionBERT, AlphaFold 3 and SPOT-RNA-1D. Molecules are either natural RNAs with homolog(s) (Natural (H)), natural RNAs without homolog(s) (Natural (nH)) or synthetic RNAs. The number of times each model outperforms the others is described in parentheses.

| Type | RNA-TorsionBERT | AlphaFold 3 | SPOT-RNA-1D |
| --- | --- | --- | --- |
| Natural (H) | **20.6 (3/5)** | 22.3 (1/5) | 22.3 (1/5) |
| Natural (nH) | **11.8 (3/3)** | 15.5 (0/3) | 13.6 (0/3) |
| Synthetic | 16.1 (2/4) | **15.9 (2/4)** | 19.1 (0/4) |
| All | **16.9 (8/12)** | 18.5 (3/12) | 18.8 (1/12) |

### Model quality assessment based on torsional angles

In this part, we describe the different datasets used for evaluating our scoring function. We used correlation scores to compare the links of our scoring function to existing metrics.

#### Datasets

Datasets of near-native structures (or decoys) are necessary to compare model quality assessment metrics. Indeed, scoring functions are used to discriminate between near-native structures, meaning that we need to have non-native structures to evaluate the quality of our scoring function.

We used three different datasets with different strategies of structure generation:

**Decoy Test Set I** is from RASP (15), composed of 85 native RNAs with decoys generated with a predictive model (by applying different sets of Gaussian restraint parameters). Each RNA has 500 decoys, which are close to the native structure. We only kept 83 RNAs and removed the two RNAs that have sequence lengths higher than 512 nucleotides (PDB ID: 3df3A and 3f1hA).

**Decoy Test Set II** corresponds to the prediction-models (PM) subset from rsRNASP (17). It has 20 non-redundant single-stranded RNAs. For each RNA, 40 decoys are generated with four RNA 3D structure prediction models The decoys are not as near to native structures as with the Decoy Test Set I. **Decoy Test Set III** is the RNA-Puzzles standardized dataset (41). This dataset comprises 21 RNAs and dozens of decoy structures for each RNA. The decoys are not all close to the native structures.

#### Evaluation measures

Scoring functions aim to discriminate near-native structures. The Pearson correlation coefficient (PCC) and the enrichment score (ES) are used to assess the correctness of a given scoring function. They assess the link between a scoring function and a given metric.

The Pearson coefficient correlation (PCC) is computed between the ranked structures based on scoring functions and structures ranked by metrics. It is defined as:

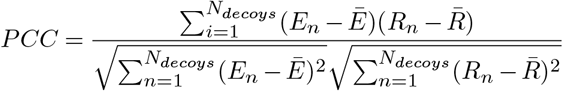

where *E*_*n*_ and *R*_*n*_ the energy and metric of the *n*th structure, respectively. PCC ranges from 0 to 1, where a PCC of 1 means the relationship between metric and energy is completely linear.

The enrichment score (ES) considers top-ranked structures from both scoring function and metric. It is defined as:

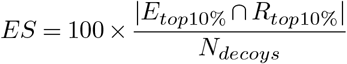

where *Etop*10% *⋂Rtop*10% is the number of common structures from the top 10% of structures (measured by the metric) and the top 10% of structures with the lowest scoring function. ES ranges between 0 and 10 (perfect scoring). An enrichment score of 1 means a random prediction, whereas below 1 means a bad score.

#### TB-MCQ as scoring function

To assess the validity of our scoring function, we computed with RNAdvisor (42) the available scoring functions RASP (15), *ϵ*SCORE (19), DFIRE-RNA (16) and rsRNASP (17) for the three different Decoys test sets. We compared TB-MCQ with the state-of-the-art scoring functions using PCC and ES with the MCQ. The averaged values are shown in Figure 6.

**Fig. 6.**
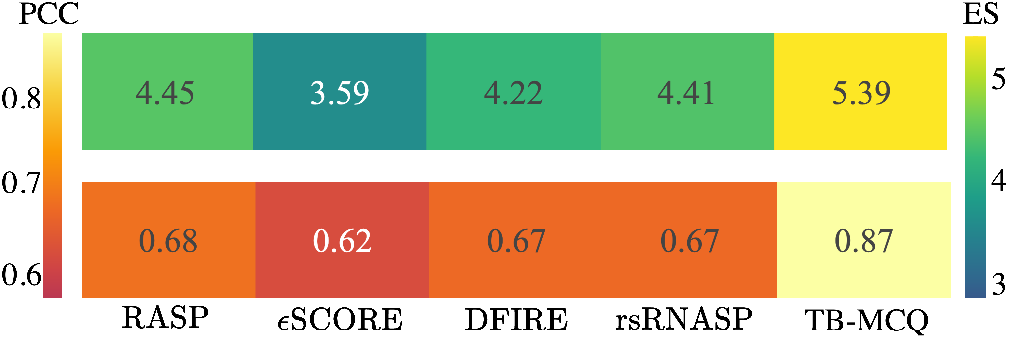
PCC and ES between five different scoring functions (RASP, ϵSCORE, DIFRE-RNA, rsRNASP and our scoring function TB-MCQ) and the angle-based metric MCQ. Values are averaged over the three decoy test sets.

TB-MCQ is the scoring function that is the more correlated to MCQ (PCC of 0.87 and ES of 5.39). rsRNASP still shows a high correlation to MCQ (PCC of 0.67 and ES of 4.41), which is surprising as it does not integrate explicit torsional angles in its computation. What is missing for both scoring functions to reproduce the MCQ metric perfectly is the accuracy of predicted torsional angles. It might be ineffective for structures that are really close to the native one and where the inferred angles from these structures are closer to the native than the predicted ones from RNA-TorsionBERT. PCC and ES for other distance-based metrics are shown in Figure S4 of the Supplementary file.

## Conclusion

In this work, we have developed a language-based model, RNA-TorsionBERT, to predict RNA torsional and pseudotorsional angles from the sequence. With a DNABERT 3-mers model, the learned embeddings have been used as a starting point to infer structural features from the sequence. We have achieved improvement compared to SPOT-RNA-1D (11), the only tool for RNA torsional angle prediction from the raw sequence.

Through an extensive benchmark of state-of-the-art methods, we have outperformed the angles inferred from the predictive models. We have also included in the benchmark the new release of AlphaFold, named AlphaFold 3 (5), which gives the best results compared to *ab initio*, template-based and deep learning solutions in terms of MCQ on inferred angles. Our method, RNA-TorsionBERT, remains better for the prediction of RNA torsional angles with only the sequence as input, while AlphaFold 3 uses MSA as inputs.

Most protein methods or current deep learning methods for predicting RNA 3D structures use MSA as inputs, which is a huge restriction. Indeed, significant families are still unknown (21). It also increases the inference time, where a homology search should be made for each prediction. Our method leverages language model without the need of homology, which is a benefit for the prediction of RNA from unknown families.

Through the evaluation of our model for backbone torsional angles prediction, we have extended this evaluation as a model quality assessment for RNA 3D structures. Then, we have inferred a scoring function named TB-MCQ. This scoring function could help the selection of near-native structures in terms of angle deviation. It is also specific to torsional angles and, thus, is more related to the angle-based metric MCQ.

Improvements could be made for both RNA-TorsionBERT and TB-MCQ. The RNA-TorsionBERT performances remain limited to reconstruct the structures from just the torsional angles. MCQ remains of high values for the different test sets, meaning there are still improvements to be made to torsional angle prediction. Indeed, the reconstruction from torsional angles alone is difficult as small angle deviation could lead to high cumulative divergence. The number of solved structures remain the main bottleneck to train robust methods. Different structural tasks could be added to the model, with the prediction of secondary structure, interatomic distances, hydrogen bonds or non-canonical base interactions. Efforts could be made to improve the language-based model used, where a model pre-trained more efficiently on RNA data could help improve the overall performances. The quality of the scoring function could be enhanced by incorporating distance atom features, or directly by improving the prediction of torsional angles itself.

Our RNA-TorsionBERT method can nonetheless be used as a starting point for the reconstruction of RNA 3D structures, with *ab initio* methods, for instance, that include molecular dynamics to relax the structure. It could also be used as a feature in a bigger network to predict RNA 3D conformation.

