## Supplementary file for "RNA-TorsionBERT: leveraging language models for RNA 3D torsion angles prediction"

December 16, 2024

### Datasets

Figure S1 shows the distribution of the ribose sugar ring angles for the different datasets used. Their distributions seem quite close, which is also the case for the pseudrotation phase $P$ angle.

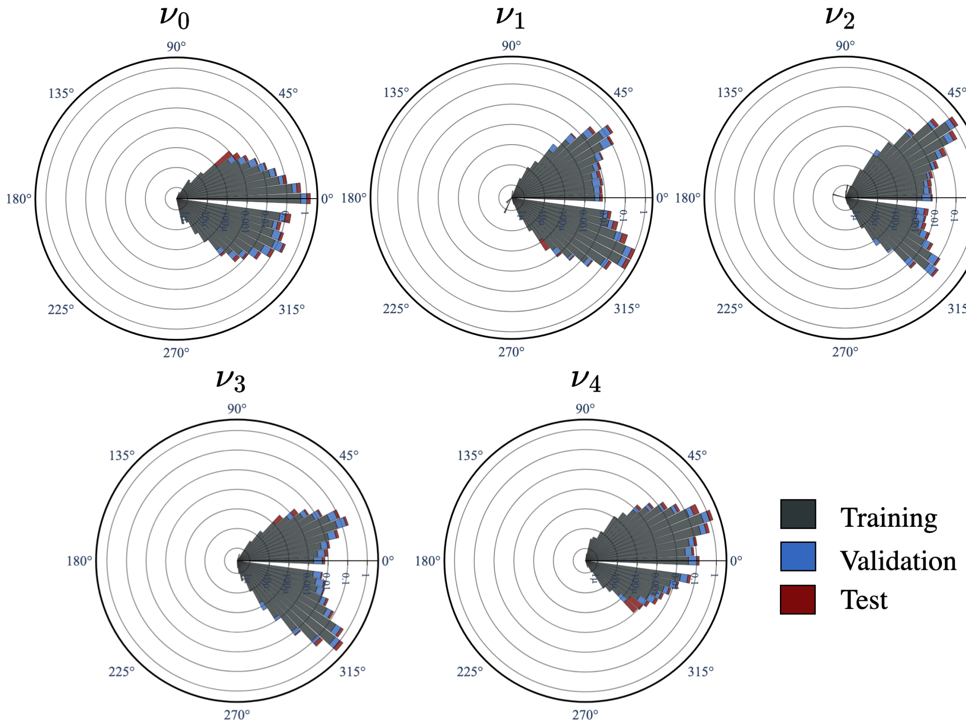


Figure S1: Polar distribution of the five ribose sugar ring angles ($\nu_0$, $\nu_1$, $\nu_2$, $\nu_3$ and $\nu_4$) for the Training, Validation and Test datasets. For each angle, the logarithm of the normalized count is depicted

### Experimental protocol

We have fine-tuned both DNABERT [1] and RNABERT [2] for the prediction of torsional and pseudo-torsional angles. For the two models, we used a batch size of 10, the Mean Average loss with a learning rate of 1e-4 and a weight decay of 0.01. We used the AdamW [3] optimizer. All inputs were padded to have a fixed size of 512 for DNABERT and 440 for RNABERT (limited by the model), and we trained the models for a maximum of 20 epochs. As there is no RNA of sequence length between 440 and 512, we used the same datasets for both RNABERT and DNABERT.

### Performances

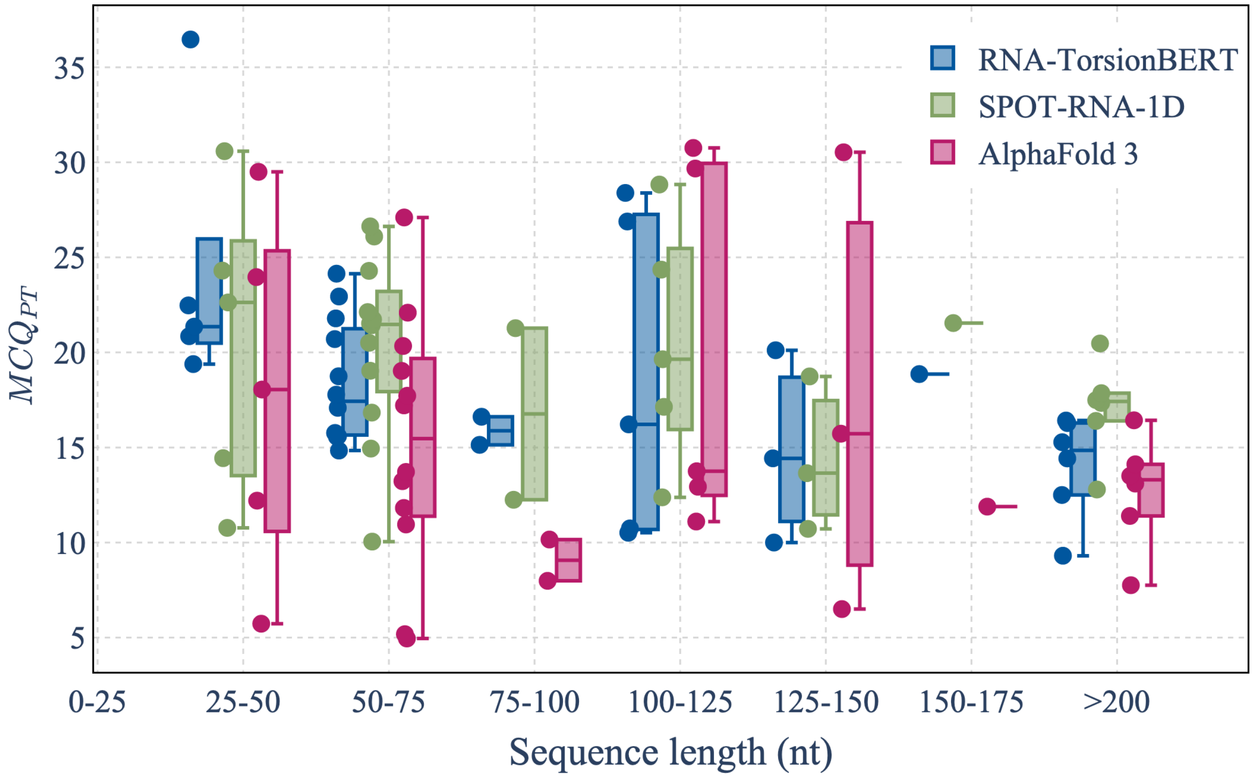


Figure S2: $MCQ_{PT}$ per window of 25nt (from 25nt to 200nt) for RNA-TorsionBERT, SPOT-RNA-1D and AlphaFold 3 inferred angles for the Test set.

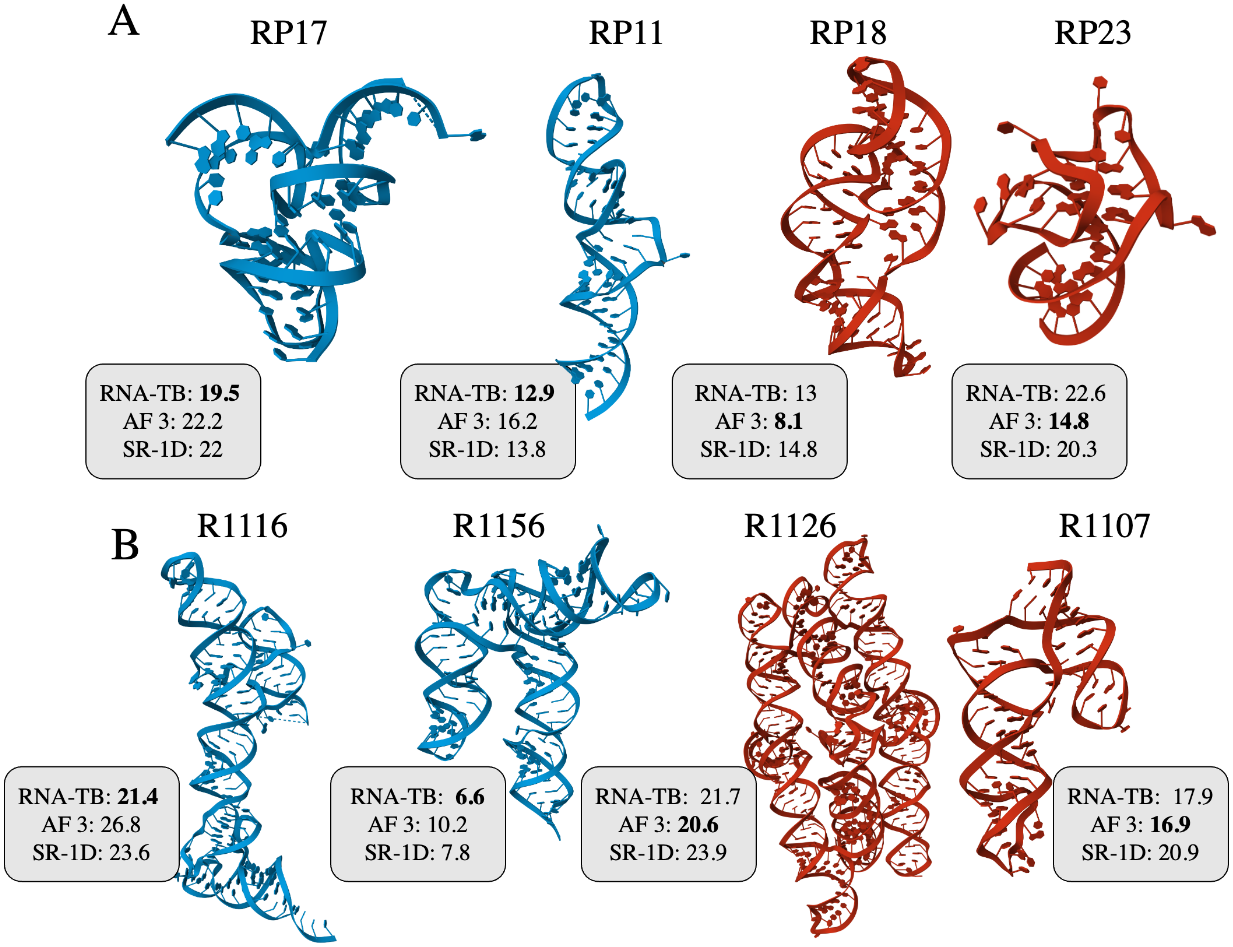


Figure S3: Structures with the associated MCQ for RNA-Puzzles (A) and CASP-RNA (B). In blue are reported examples of structures where RNA-TorsionBERT outperforms AlphaFold 3 and SPOT-RNA-1D. In red are examples of RNA structures where AlphaFold 3 outperforms RNA-TorsionBERT and SPOT-RNA-1D.

Table S1: MCQ per pseudo-torsional angle and $MCQ_{PT}$ (MCQ computed for all the pseudo-torsional angles) over the Test set for RNA-TorsionBERT compared to SPOT-RNA-1D [4]. We also include inferred torsional angles from state-of-the-art methods that predict RNA 3D structures from State-of-the-RNArt [5]. Methods are sorted by $MCQ_{PT}$.

| Models | $MCQ_{(\eta)}$ | $MCQ_{(\theta)}$ | $MCQ_{PT}$ |
|---|---|---|---|
| RNA-TorsionBERT | 15.2 | 20.8 | 18.0 |
| SPOT-RNA-1D [4] | 17.0 | 21.3 | 19.1 |
| AlphaFold3 [6] | **13.8** | **17.4** | **15.6** |
| IsRNA1 [7] | 18.9 | 26.1 | 22.4 |
| RNAJP [8] | 20.3 | 25.5 | 22.8 |
| Vfold-Pipeline [9] | 21.0 | 27.6 | 24.2 |
| RNAComposer [10] | 21.0 | 28.1 | 24.5 |
| 3dRNA [11] | 25.5 | 31.6 | 28.5 |
| RhoFold [12] | 28.1 | 31.6 | 29.8 |
| MC-Sym [13] | 28.5 | 32.9 | 30.6 |
| trRosettaRNA [14] | 26.0 | 36.9 | 31.3 |

Table S2: $MCQ_{PT}$ for our method RNA-TorsionBERT, AlphaFold 3 [6] and SPOT-RNA-1D [4] on secondary motifs averaged on the Test set. Secondary motifs are extracted from RNApdbee [15]

| Motifs | RNA-TorsionBERT | AlphaFold 3 | SPOT-RNA-1D |
|---|---|---|---|
| Single-stranded | 36.4 | **31.9** | 48.4 |
| Loops | 31.3 | **30** | 42.3 |
| Stems | 16.3 | **15.6** | 24.2 |

Table S3: MCQ per RNA family for the single-stranded structures from RNA-Puzzles [16–19] dataset. The number of times each model outperforms the others is described in parentheses. The models compared are RNA-TorsionBERT, AlphaFold 3 [6] and SPOT-RNA-1D [4].

| Family | RNA-TorsionBERT | AlphaFold 3 | SPOT-RNA-1D |
|---|---|---|---|
| Aptamer | 18.6 (0/3) | **16.4** (**2**/3) | 17.5 (1/3) |
| Riboregulator | **13.0** (**1**/1) | 16.2 (0/1) | 13.8 (0/1) |
| Riboswitch | 16 (1/11) | **13.9** (**9**/11) | 16.6 (1/11) |
| Ribozyme | **22.6** (**3**/4) | 23.0 (1/4) | 24.5 (0/4) |
| Ricin loop | 8.5 (0/1) | **6.6** (**1**/1) | 10.9 (0/1) |
| Virus | 13.1 (0/2) | **10.3** (**2**/2) | 16.5 (0/2) |
| All | 16.8 (5/22) | **15.3** (**15/22**) | 17.7 (1/22) |

### Model quality assessment based on torsional angles

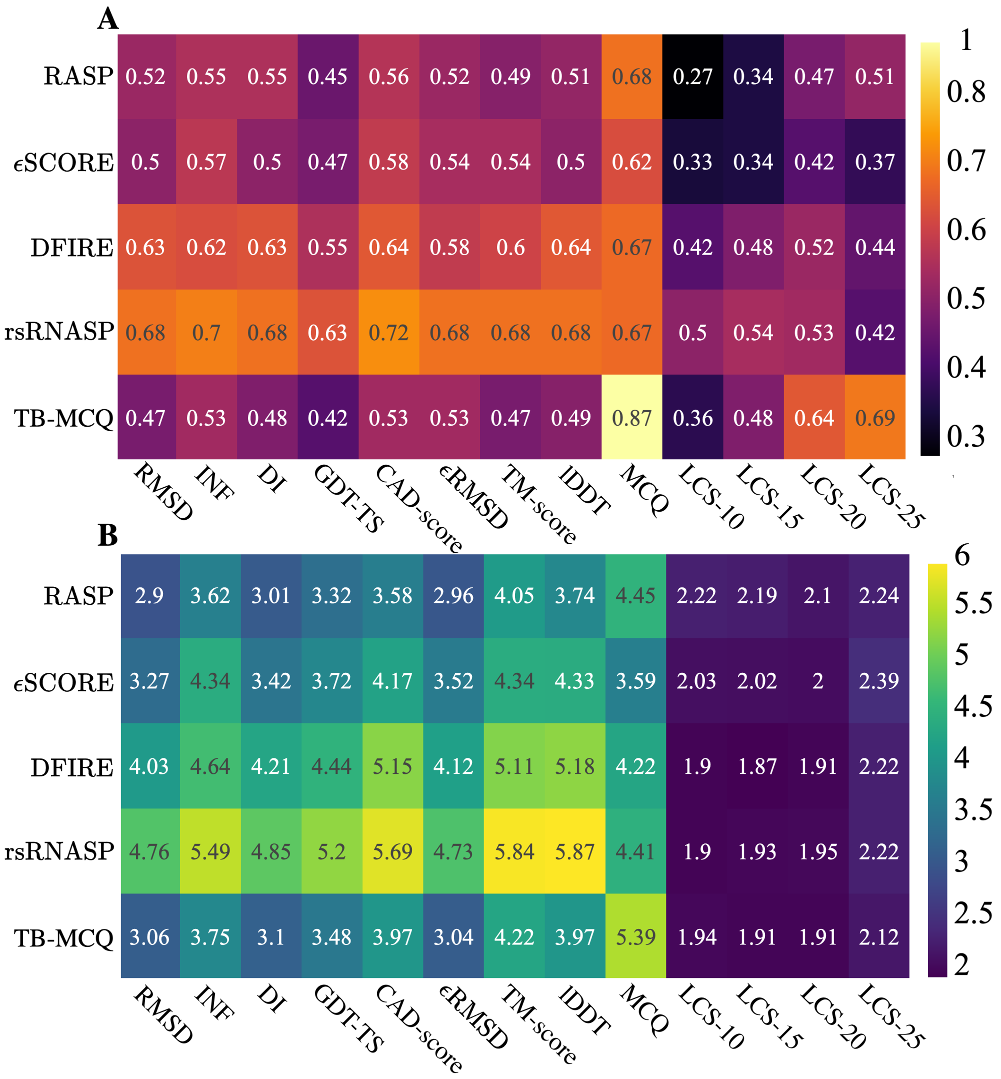


Figure S4: PCC (A) and ES (B) between five different scoring functions (RASP [20], $\epsilon$SCORE [21], DIFRE-RNA [22], rsRNASP [23] and our scoring functions TB-MCQ) and ten metrics (RMSD, $\text{INF}_{all}$ [24], DI [24], GDT-TS [25], CAD-score [26], $\epsilon$RMSD [21], TM-score [27, 28], lDDT [29], MCQ [30], and LCS-TA [31] (with a threshold of 10, 15, 20 and 25)). Values are averaged over the three decoy test sets.
